# Are resting-state networks the brain’s cognitive atoms? Differential dynamic reconfiguration of functional brain networks across tasks and at rest

**DOI:** 10.1101/2024.11.06.622287

**Authors:** JeYoung Jung, Matthew A. Lambon Ralph

## Abstract

It is increasingly popular to utilise functional connectivity (FC) analyses of resting-state functional magnetic resonance imaging (fMRI) to characterize human functional brain networks and to use the emergent resting-state networks (RSNs) in basic and clinical neuroscience. Often, they are treated as ‘atomic’ building blocks that underpin human cognition. However, the true function of these RSNs, as well as the relationship between intrinsic and task-evoked functional brain networks, are complex and incompletely characterized. Here, we investigated the functional characteristics of the intrinsic and extrinsic networks using resting-state and task fMRI. Independent component analysis (ICA) was used to estimate spatiotemporal functional networks during tasks and at rest, and to compare the spatiotemporal properties of each network. While there was some spatial correspondence between the RSNs and task-evoked networks, our results demonstrated that the task-evoked functional networks were different from the RSNs in task-relatedness as well as spatial topology. Furthermore, the degree of topological differences between the RSNs and task-evoked networks was modulated by a given task. Comparison between the RSNs and task-evoked networks showed that tasks reconfigure the RSNs by changing FC with various brain regions specific to the task condition. Our findings indicate that the brain does not maintain an “invariant intrinsic” network architecture when it engages in a task. Instead, the tasks reconfigure the network architectures, thereby accommodating specific computational/representational task requirements through flexible interactions between demand-specific regions. Thus, the results suggest that task fMRI is required to understand the full repertoire of the brain’s functional architecture.

**Significant Statement:** Resting-state networks (RSNs) could offer a critical foundation for understanding the brain’s intrinsic organization. However, the functional nature of these intrinsic networks, and their relationship to those activated during specific tasks, are complex and not fully understood. We undertook a comprehensive examination of the functional attributes of intrinsic and extrinsic networks. Although we observed some spatial congruence between RSNs and task-evoked networks, we found fundamental differences in their task-relatedness and spatial topology. These findings highlight the dynamic nature of the brain’s functional networks, which adapt to specific task demands through flexible interactions among task-specific regions. Thus, task fMRI is essential for a comprehensive understanding of how the human brain dynamically reconfigures its functional architecture in response to external demands, providing valuable insights into the cognition and behaviour.

## Introduction

The human brain supports a huge array and variety of mental and physical activities. It seems very unlikely that the brain achieves this by “neuromarquetry” with each task supported by a separate brain area (there are too many tasks, and we are capable of undertaking entirely novel tasks), instead it seems more likely that it is achieved by dynamically combining more generalised neurocognitive components according to the task requirements ^1, 2^. Much like molecular chemistry, cognitive activities (molecules) arise from different combinations of ‘cognitive atoms’ or even of the same atoms (e.g., oxygen and water combining to make water or hydrogen peroxide: H_2_O vs H_2_O_2_). Recently, an influential notion has suggested that resting-state functional connectivity (FC) can characterize the brain’s intrinsic functional network architecture ^3, 4^. Are these emergent resting-state networks (RSNs) “cognitive atoms”? If so, like Dalton’s atoms, they should be indivisible and invariant and have describable properties.

Initial studies examined task-related FC to map functional brain networks ^5, 6^. Then Biswal et al ^7^ demonstrated that, during rest, the left and right primary motor cortices were strongly correlated in their resting-state blood-oxygen-level dependent (BOLD) time series: i.e., the primary motor network (MN; which overlaps with the task-active motor system). Numerous studies have since replicated these pioneering findings and extended to other parts of the brain, revealing other relatively stable RSNs such as the primary visual network (VN1), auditory network and higher order cognitive networks ^3, 4, 8, 9, 10^. These RSNs consist of anatomically separated but linked brain regions that show strong FC at rest and partially overlap with functional networks derived from task-fMRI studies, providing clues about the potential functional relevance of the RSNs. In addition, the practical convenience of the technique (relatively short scanning, 5-10mins, without a task) has resulted in an explosion of research exploring functional alterations in the RSNs in neurological and psychiatric brain disorders ^11, 12, 13, 14^.

To link the resting-state FC to cognition, a relatively small number of studies have probed for the correspondence between intrinsic RSNs derived from rsfMRI data and task-evoked coactivation FC using task-fMRI (tfMRI) with various tasks ^9, 15, 16^, via large-scale meta-analytic approaches ^17, 18^ or various other methods (for the review, see ^4^). Employing open source big database (e.g., Human Connectome Project: HCP) with rsfMRI and various tfMRI (7 tasks), Cole et al ^15^ suggested that there is an intrinsic network architecture present across many tasks and it shapes functional network architecture during a given task. However, recent studies have emphasized sometimes considerable differences in FC patterns during task and at rest ^19, 20, 21^. Thus, some studies support a universal architecture, whilst others defend differential resting and task network architectures.

Here, we investigated the functional characteristics of the intrinsic and extrinsic networks using rsfMRI and tfMRI. To provide a formal and detailed comparison between them, we used a data-driven multivariate approach - independent component analysis (ICA) to estimate spatiotemporal functional networks during tasks and at rest. We conducted three ICAs for rsfMRI, tfMRI and the combined data (rsfMRI + tfMRI) to define brain networks and compared the properties of networks derived from the three ICAs. If the architecture of resting and task-evoked networks were similar, then the networks estimated the combined ICA would produce similar network profiles related to task states – task-relatedness from tfMRI. Then, we computed the spatial similarity between networks estimated from the three ICAs to identify the matched networks and compared the degree of similarity between them. Furthermore, to evaluate the characteristics of networks, we directly compared the matched networks from tfMRI and rsfMRI – if they were similar to each other, there would be null result in this comparison. To ensure generalizability, we applied this approach to the open-source data from HCP including rsfMRI and three tfMRI (language, working memory and motor tasks).

## Materials and Methods

### Participants

We recruited twenty-one healthy English native speakers (7 males, mean age = 23 years ± 4, range from 20 to 35 years). All participants were right-handed as assessed by the Edinburgh Inventory for Handedness ^22^. After a detailed explanation of the study, all participants gave their written informed consent. The experiment was approved by the ethics committee of the University of Manchester in accordance with the Declaration of Helsinki.

To replicate and extend our findings, we also used the open source data from Human Connectome Project (HCP) ^23^. Participants were recruited from Washington University and gave informed consent approved by the Washington University Institutional Review Board. The data used were from the “1200 Subjects” HCP Young Adult release. Twenty-one subjects (7 males, age ranging from 20 to 35 years) from this data set completed task-fMRI were used, matching age and gender with our data.

### Experimental procedures and task paradigms

Our data were collected using rsfMRI and tfMRI. rsfMRI (6mins, eyes open with fixation) was collected prior to tfMRI (10mins). During tfMRI, participants performed a semantic association task and a pattern matching task (control task). Participants saw three pictures on the screen (top: target, bottom: choices) in each trial. The semantic task asked participants to decide which of two bottom pictures (e.g., grapes – related item, orange – unrelated item) was more related in meaning to a target top picture (e.g., wine). The items for the semantic association task were created by combining the Pyramids and Palm Trees test (PPT) ^24^ and an abridged version of the Camel and Cactus test (CCT) ^25^. The pattern matching task asked participants to choose which of two patterns was identical to a target pattern. The items for the pattern matching task were created by scrambling the pictures used in the semantic task. There were 9 blocks of the semantic and control task interleaved (e.g., A-B-A-B). A task block had 4 trials, and a trial started with 0.5s fixation followed by the items presented for 4.5s. Fixation blocks (4s) were placed between the task blocks. Participants pressed one of two buttons designating two choices in a trial. E-prime software (Psychology Software Tools Inc., Pittsburgh, USA) was used to display stimuli and to record responses.

HCP data were collected over 2 days. On each day, rsfMRI ^26^ was acquired with 2 runs (eyes open with fixation, 28mins for a run and 56mins total) followed by 30mins of task-fMRI (60mins total) ^27^. During tfMRI, participants performed a set of seven tasks including emotion, reward learning, language, motor, relational reasoning, social cognition and working memory. Here, we chose 3 tasks for the analysis: language, motor and working memory (WM). The language task used auditory stimuli consisting of narrative stories and math problems, along with questions to be answered regarding the prior auditory stimuli. The motor task asked participants to move the hands tongue and feet following the visual instruction. The WM task involved a visual N-back task (2-back vs. 0-back), in which participants indicate a match of the current item to either a constant target item or two items previous. For the detailed description of each task, see the Barch et al (2013) ^27^.

### fMRI data acquisition and analysis

Whole brain imaging was acquired with a 3T Philips Achieva scanner using a 32-channel head coil with a SENSE factor 2.5. We utilised a dual-echo fMRI protocol developed by Halai et al ^28^ to maximise signal-to-noise (SNR) in the orbitofrontal regions and anterior temporal areas. fMRI parameters included 42 slices, 96 x 96 matrix, 240 x 240 x 126mm FOV, in-plane resolution 2.5 x 2.5, slice thickness 3mm, TR = 2.8s, TE = 12ms and 35ms. A T1-weighted structural image was acquired using a 3D MPRAGE pulse sequence with 200 slices, in-planed resolution 0.94 x 0.94mm slice thickness 0.9mm, TR = 8.4ms, TE =3.9ms.

Image processing was carried out using MATLAB R2012a and SPM8. The dual gradient echo images were extracted and averaged using in-house MATLAB code developed by Halai et al ^28^. Functional images were realigned correcting for motion artefacts and different signal acquisition times by shifting the signal measured in each slice relative to the acquisition of the middle slice prior to combining the short and long echo images. The mean functional EPI image was co-registered to the individual T1-weighted image and segmented using the DARTEL (diffeomorphic anatomical registration through an exponentiated lie algebra) toolbox ^29^. Then, normalization was performed using DARTEL to warp and reslice images into MNI space and smoothing was applied with an 8mm full-width half-maximum Gaussian filter.

HCP data were acquired with a 32-channel head coil on a modified 3T Siemens Skyra with TR = 720 ms, TE = 33.1 ms, flip angle = 52°, BW = 2,290 Hz/Px, in-plane FOV = 208 × 180 mm, 72 slices, 2.0 mm isotropic voxels, with a multiband acceleration factor of 8 ^30^. Preprocessing was performed by the HCP data analysis pipelines using FSL ^31^. It includes gradient unwarping, motion correction, fieldmap-based EPI distortion correction, brain-boundary-based registration of EPI to structural T1-weighted scan, non-linear (FNIRT) registration into MNI152 space, and grand-mean intensity normalization. For the comparison with our dataset, we applied an 8mm full-width half-maximum Gaussian filter for smoothing.

SPM12 was used to perform a general linear model (GLM). At the individual level, a design matrix was defined by modelling task conditions, according to each dataset. Additionally, six motion parameters were included as regressors in the design matrix. In our dataset, semantic and control conditions were modelled as the contrasts of interest. In HCP dataset, task conditions were entered as the contrast of interest for each tfMRI: language – story and math; motor – movement; WM – 2-back and 0-back. Contrast images were computed to assess difference in activation in each task condition for each participant. Group analysis was conducted using a random effect model (one-sample t-test). Statistical threshold was set at p < 0.001 at the voxel-level and p < 0.05 at the cluster level with at least 100 contiguous voxels after family-wise error (FWE) correction.

### Independent component analysis

Independent component analysis (ICA) is a data-driven multivariate approach to decompose a mixed signal into independent components (ICs) ^32^. ICA utilizes fluctuation in the fMRI data to separate the signal into maximally independent spatial maps (components), each explaining unique variance of the 4D fMRI data. Each component has a time course related to a coherent neural signalling associated with a specific task, artifact, or both.

ICA was performed using a group ICA algorithm (GIFT, http://icatb.sourceforge.net/, version 3.0a) ^33^. Preprocessed images were entered the toolbox. Using Maximum Description Length (MDL) and Akaike’s criteria, the number of independent components was estimated. A first stage subject-specific principal components analysis (PCA) was performed. A second stage group data reduction, using the expectation-maximization algorithm included in GIFT. Then Informax ICA algorithm ^34^ was conducted, repeating it 20 times in ICASSO implemented in GIFT to generate a stable set of final components. Finally, the ICs were then estimated using the GICA back-reconstruction method based on PCA compression and projection ^32^.

In order to compare resting and task-related networks, we conducted ICAs for the combined rsfMRI with tfMRI, rsfMRI and tfMRI separately in each dataset (Fig. 1A). The resultant ICs were inspected visually to exclude residual artefact such as the signal around the edge of brain and within cerebral spinal fluid spaces. The remaining ICs were defined as ‘networks’. We labelled them with regional or functional descriptors (e.g., default mode network; DMN, motor network; MN). Then, we investigated if ICs were significantly involved in any task condition using the temporal sorting in GIFT. Temporal sorting was conducted by applying a GLM to the component’s time course (Fig. 1B). The fMRI run specific time course for each subject were regressed against the design matrix for the tasks and tested for significance to identify components where activity was greater during each task condition against rest. The resulting beta (β) weights represent the degree to which component network recruitment was modulated by the task conditions. For a given component, positive and negative β weights indicate task-related network recruitment that is increased or decreased with respect to baseline, respectively.

**Figure 1.**
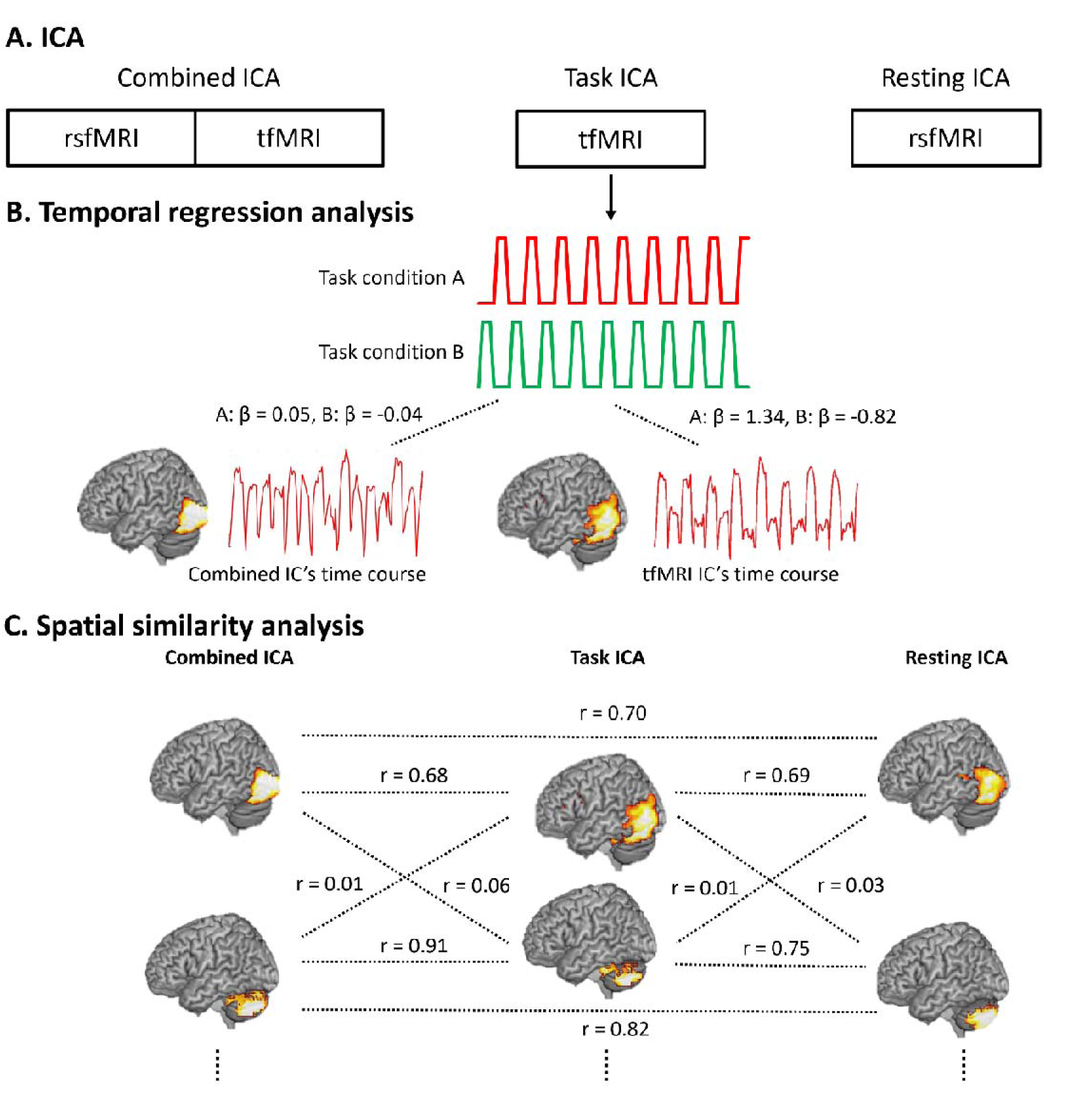
A) The procedures of ICA. B) Temporal regression in the combined and task ICA. C) Spatial correlation between the networks estimated from three ICAs.

Only tfMRI session from the combined and task ICAs was used for this analysis because there was no task during rsfMRI. To evaluate the task-related network recruitment, we performed a one-sample t-test on the β weights (p _FDR-corrected_ < 0.05).

To find the matched ICs from 3 ICAs (the combined, task and resting ICAs), we evaluate the spatial similarity between the components using the spatial sorting in GIFT (Fig. 1C). Spatial sorting was conducted by correlating the group ICs’ spatial maps with the spatial map of the other ICA’s group mean IC as a template. The ICs’ spatial maps were thresholded at p < 0.001 at the voxel-level and p < 0.05 at the cluster level with at least 100 contiguous voxels after family-wise error (FWE) correction. The resulting correlation coefficients represent the degree to which IC was spatially overlapping with the IC from the other ICAs’ ICs. An IC showed the highest correlation coefficient was defined as the matched component. Then, we performed paired t-tests on the matched ICs between Task and Resting ICAs using SPM12 to investigate the functional characteristics of the network during task and at rest. Finally, to evaluate the spatial similarity, we performed a one-sample t-test on the correlation coefficients (p _FDR-corrected_ < 0.05). For the comparison of spatial similarity between different ICAs, paired t-tests were performed for each component from HCP dataset (p _FDR-corrected_ < 0.05).

## Results

### GLM results

Whole brain analysis revealed that the semantic task evoked significant activation in bilateral inferior frontal gyrus (IFG), anterior temporal lobe (ATL), fusiform gyrus, posterior middle temporal gyrus (pMTG) and middle frontal gyrus and the control task induced the significant activation in bilateral superior parietal lobe (SPL), inferior parietal lobe (IPL), precuneus and superior frontal gyrus (SFG) (Fig. 2A).

**Figure 2.**
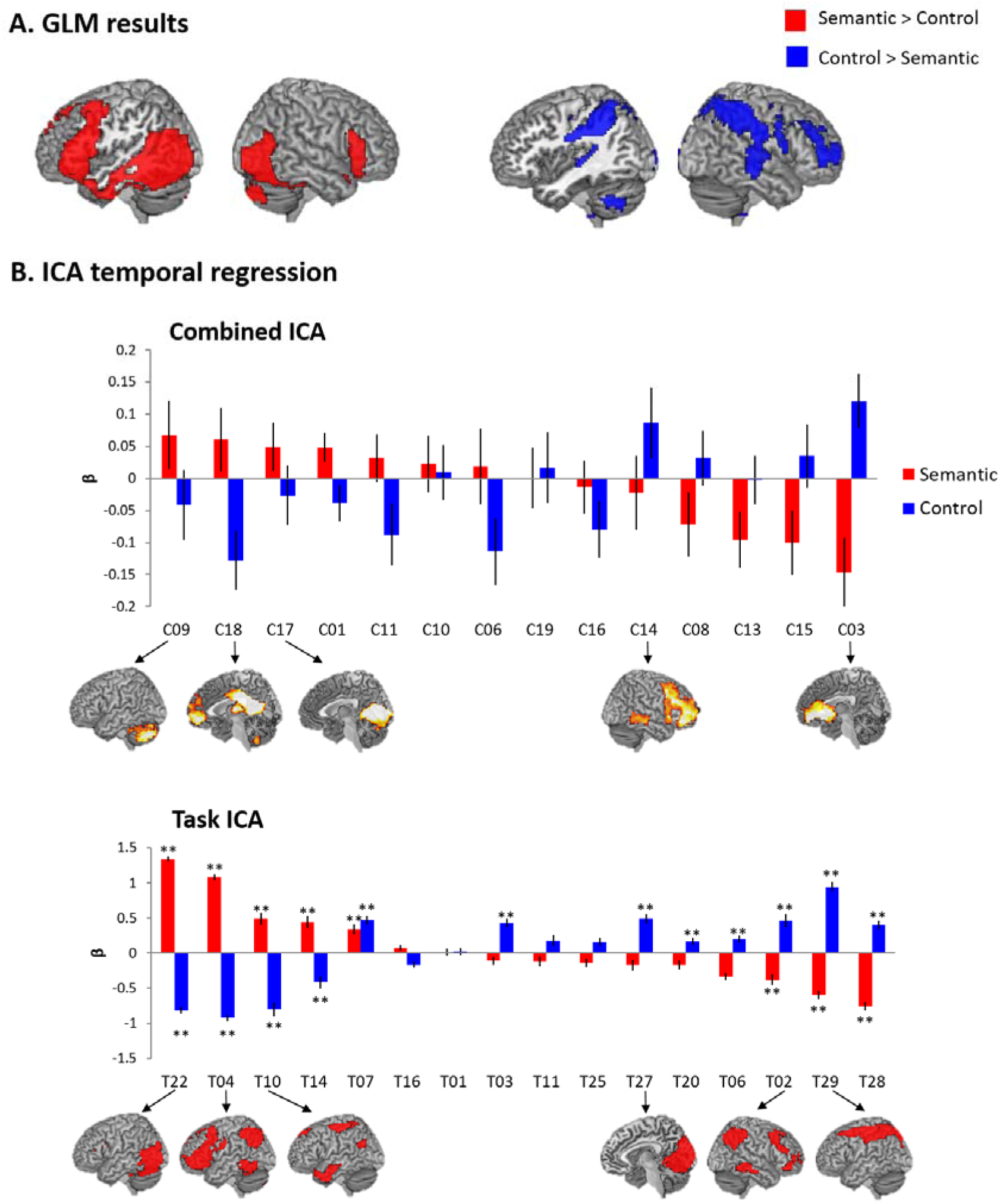
A) The results of GLM in our dataset (semantic). B) The results of ICA temporal regression. Top: the temporal regression of the combined ICs, Bottom: the temporal regression of the task ICs. ** p _FDR-corrected_ < 0.005

The GLM results of HCP data are summarised in Supplementary Info (Fig. S1). In the HCP language data, the story condition evoked significant activation in the bilateral IFG, ATL, superior middle temporal gyrus (STG), MTG, medial prefrontal gyrus (mPFC), dorsal mPFC, left angular gyrus (AG) and middle cingulate cortex (MCC). The math condition increased activation in the bilateral dorsolateral prefrontal cortex (DLPFC), IPL, insular, and MCC. The HCP motor data showed the activation in bilateral pre/postcentral gyrus, supplementary motor area (SMA), rolandic operculum (RO), cerebellum and left supuramarginal gyrus during the motor task. The working memory data showed activation in the bilateral DLPFC, insular, SMA, anterior cingulate cortex (ACC), inferior parietal sulcus (IPS), precuneus, visual cortex and cerebellum. These results replicate the original findings from Barch et al (2013) ^27^.

### ICA results – temporal regression

We found 26 ICs in the combined ICA (12 noise ICs), 32 ICs in task ICA (18 noise ICs) and 26 ICs in rest ICA (11 noise ICs) in our dataset (Fig. S2). In the HCP dataset, the combined ICAs had 20 ICs each in the combined language (10 noise ICs), motor (8 noise ICs) and WM (8 noise ICs), the task ICAs resulted in 16 ICs for language (6 noise ICs), 17 ICs for motor (7 noise ICs) and 16 ICs for WM (9 noise ICs), and the rest ICA showed 25 ICs (11 noise ICs) (Fig. S3-S6).

First, we examined the involvement of task conditions with ICs from the combined ICA and task ICA to compare the characteristics of brain networks derived from rest and task fMRI. If there were no differences between resting and task-related brain networks, the results of task-relatedness in each ICA would be similar. For example, the semantic condition significantly activated the semantic, cognitive control and visual networks in both the combined ICA and task ICA (Fig. 2A). However, we found that the ICs from the combined ICA did not show any significant involvement in either semantic or control (pattern matching) conditions (Fig. 2B). Also, the IC that showed the strongest activation during semantic processing was the cerebellum (C09), whereas the IC for the control processing was the medial prefrontal cortex (mPFC, C03). Contrary to the combined ICA, the task ICA mirrored the results corresponding to the GLM results. As we expected, the semantic condition evoked the significant activation in the secondary visual network (VN2: T22/T07), the left frontoparietal network (l.FPN: T04), the semantic network (SN: T10) and the default mode network (DMN: T14). The control condition was significantly associated with the superior parietal lobe (SPL: T29), the right frontoparietal network (r.FPN: T02), the primary VN (VN1: T27), the VN2 (T07), the DMN (T28), the mPFC (T20), the cerebellum (T03), and the rolandic operculum network (RON: T06) (Fig 2B & Table S1).

The results of HCP datasets also showed the discrepancy between the combined ICA and task ICA. The combined ICA with resting and language fMRI showed that the story condition activated the auditory network (C07), the SN (C06) and the mPFC (C11), whereas the math condition was associated with the FPN (C02), salience network (C04), and the dorsomedial prefrontal cortex (dmPFC: C13) in order of task relevance (Fig. S3). The language task ICA revealed that the story condition activated the SN (L02), the auditory network (L06) and the mPFC (L16), whereas the math condition induced the significant activation in the FPN (L05) and dmPFC (L08) (Fig. S3). Contrary to the combined ICA, the task ICA demonstrated differential task modulation in each network (Fig. S3 & Table S2). The involvement of the story condition was higher in the SN than the auditory network (p < 0.05) and the math condition involvement was stronger in the FPN than dmPFC (p < 0.005) – these were not found in the combined ICA results.

The combined resting and motor fMRI showed involvement of the cerebellum (C114) and the sensory-motor network (C10) during movements. The motor task ICA revealed that movement condition was associated with the cerebellum (M05), the motor network (MN: M02) and the secondary sensory network (M04) (Fig. S4).

The combined resting and WM fMRI revealed that the 2-back condition evoked activation in the VN2 (C11), the FPN (C03), the dmPFC (C08) and the cerebellum (C06) and the 0-back condition activated the VN2 (C11) (Fig. S4). The VN2 (C11) and the FPN (C03) were more involved in the 2-back condition than the 0-back condition (p < 0.005). The WM task ICA showed that the VN2 (W01), the FPN (W07), the dmPFC (W15) were significantly associated with both 2-back and 0-back conditions (Fig. S5). The cerebellum (W04) was involved in the 2-back task. Contrary to the combined ICA, the task ICA demonstrated an effect of working memory load in the related brain networks including FPN and VN2: the 2-back condition activated the FPN more strongly than the 0-back condition (p < 0.005) and activated the VN2 less than the 0-back task (p < 0.005) (Fig. S5).

### ICA results – spatial similarity

Second, we investigated the spatial similarity between ICs estimated from three ICAs (the combined, task and rest ICAs). We identified the matched ICs between the combined, task and rest ICAs and compare the spatial similarity between them. To compare the degree of spatial overlapping between the matched ICs across the datasets, we chose three task-general networks including the VN1, FPN and DMN found in all ICAs and three task-specific networks, including the SN, VN2 and MN, that showed the strongest task involvement in temporal regression from each ICA (Fig. 3A & B).

**Figure 3.**
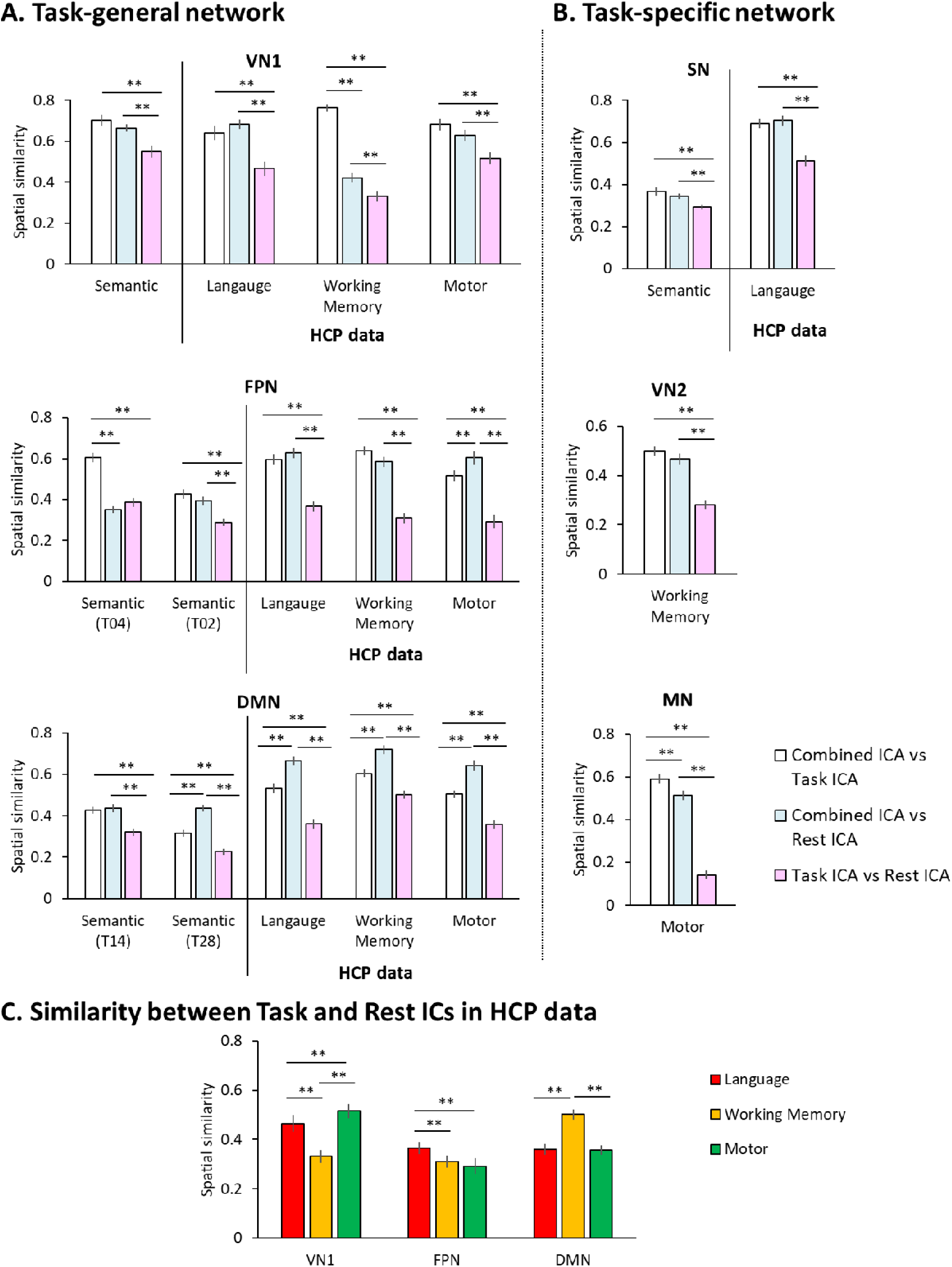
A) The results of spatial correlation in the task-general networks. B) The results of spatial correlation in the task-specific networks. White bars indicate the spatial similarity between the combined and task networks. Grey bars indicate the spatial similarity between the combined and rest networks. Pattern bars indicate the spatial similarity between the task and rest networks. C) The similarity between task and rest networks in HCP data. ** p < 0.005

The VN1 showed the significant decrease of the spatial similarity between rest and task ICA compared to the similarity of other ICAs (the combined ICA + task ICA and the combined ICA + rest ICA) in our data (semantic) and all HCP data (language, motor and WM) (p _FDR-corrected_ < 0.005). The WM data revealed that the similarity between the combined ICA and rest ICA was significantly decreased compared to that between the combined ICA and task ICA (Fig. 3A **VN1**). The spatial overlapping between rest and task ICA in the FPN was the lowest across all datasets (our data had two FPNs, the left [T04] and right FPN [T02]). The motor data also showed that the similarity between the combined ICA and rest ICA was significantly higher than the other pairs (Fig. 3A **FPN**). The DMN also showed similar results with the other task-general networks. However, the similarity between the combined ICA and rest ICA was higher than the other ones in our data (semantic [T28]) and all HCP datasets (Fig. 3A **DMN**).

Furthermore, task-specific networks in each dataset also demonstrated the similar results with task-general networks. The SN found in semantic and language fMRI revealed significantly decreased spatial overlapping between rest ICA and task ICA than any other pairs (Fig. 3B **SN**). The VN2 in WM data and MN in the Motor data also showed the same pattern of results (Fig. 3B). Overall, all networks showed significantly reduced spatial similarity between task and rest ICAs compared to the similarity observed in other ICAs (the combined ICA + task ICA and the combined ICA + rest ICA) (ps < 0.6).

If a task shapes brain networks differently from resting, the cognitive requirements of tasks should modulate the functional characteristics of networks. To examine this hypothesis, we compared the spatial similarity between task and rest ICAs in the HCP datasets (language, motor and WM). Each task modulated three task-general networks differently, resulting in significantly different spatial similarity across the three task fMRIs (Fig. 3C). The WM task (pictures of faces, places, tools and body parts) led to the biggest discrepancy between task and rest VN1 compared to language (auditory) and motor (word instruction) tasks. The FPN showed significantly reduced similarity during motor task compared to language and WM tasks. The DMN showed the highest similarity during 2-back processing (WM) compared to other tasks.

### Comparison of task and rest networks

Finally, we compared the matched ICs from task and rest ICA to examine the functional characteristics of the network during task and at rest. We present the results from the task-general tasks (VN1, FPN and DMN) and task-specific networks (SN, VN2 and MN).

Fig. 4 illustrates the results of the VN1. In our data, the VN1 (T27) was more synchronized with the calcarine gyrus, middle occipital gyrus (MOG), cuneus, posterior middle temporal gyrus (pMTG) and the cerebellum, whereas desynchronized with the calcarine gyrus, inferior occipital gyrus (IOG), precuneus, and the right middle frontal gyrus (MFG) during task fMRI (below the dashed line). In the HCP data, the language task (L01) led to co-deactivation of the VN1 with the calcarine gyrus, cuneus, superior temporal gyrus (STG), thalamus, and the precentral gyrus. The WM task (W09) evoked significant coupling of the posterior cingulate cortex (PCC) and the precuneus as well as co-deactivation in the MOG, IOG, anterior temporal cortex (ATL) and the MFG with the VN1. During the motor task, the VN1 (M01) was coupled with the cerebellum, IOG, lingual gyrus, inferior temporal gyrus (ITG), fusiform gyrus (FG), inferior parietal lobe (IPL), superior parietal lobe (SPL), precuneus, angular gyrus (AG), right superior frontal gyrus (SFG), MFG, MTG, caudate, putamen, and the mPFC, whereas decoupled with the inferior frontal gyrus (IFG) and the cuneus.

**Figure 4.**
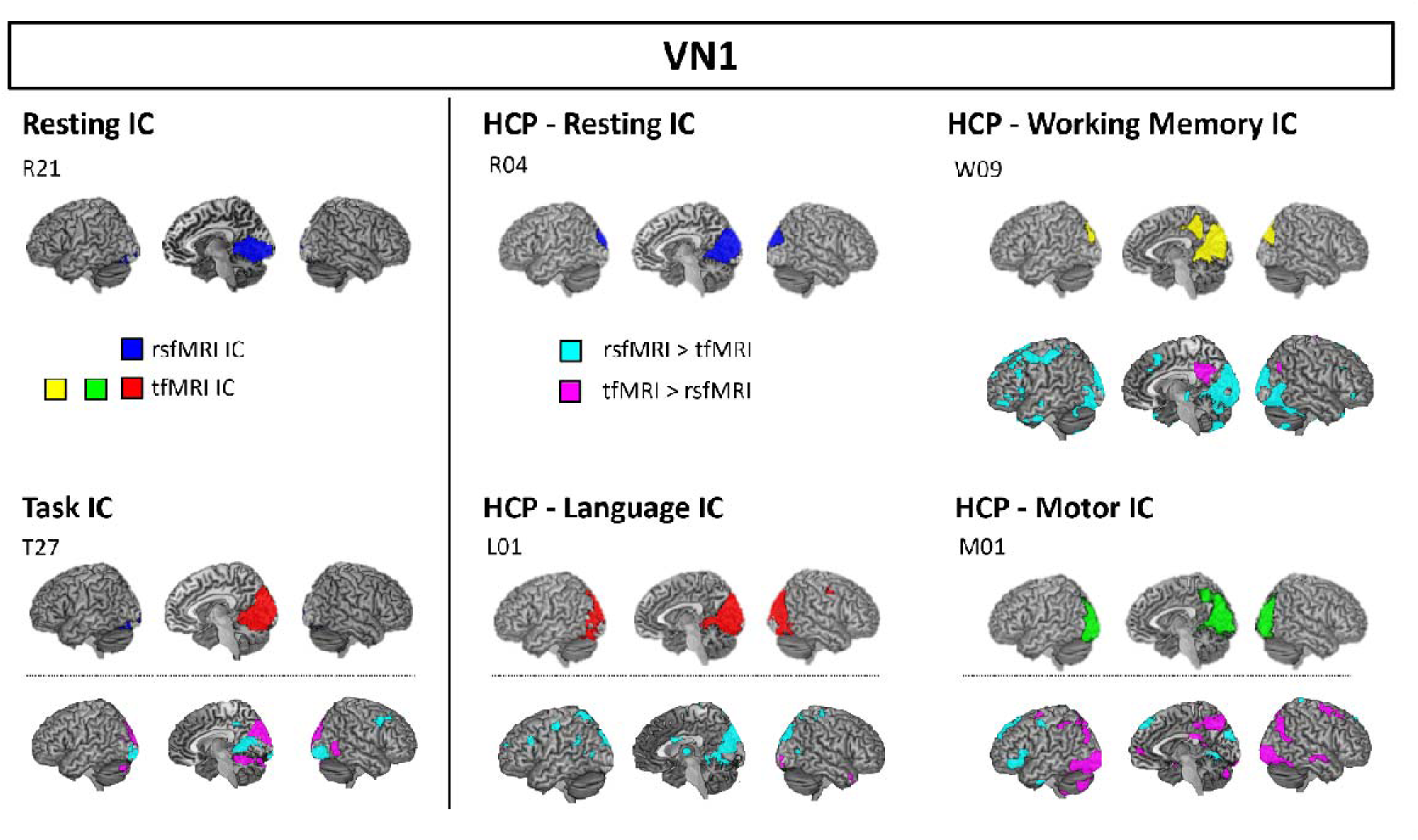
The comparison between task and rest VN1. The ICs above a dashed line are the estimated VN1 from each dataset. Under the dashed line, the results of the comparison between task and rest ICs are displayed. Cyan colour indicates the significant coupling in rsfMRI compared to tfMRI whereas pink colour indicates the significant coupling during tfMRI relative to rsfMRI.

Fig. 5 displays the results for the FPN. In our data, we had two FPNs one in each hemisphere (l.FPN and r.FPN) in the rest and task ICAs. During task fMRI, the l.FPN (T04) was significantly synchronised with the left ATL, hippocampus, bilateral IPL, SPL, AG, mPFC, anterior cingulate cortex (ACC), SFG, precentral/postcentral gyrus, PCC, calcarine gyrus, IOG and the cerebellum. During rest more than during task, the l.FPN (T04) was significantly coactivated with the precuneus, right IFG, MFG, AG, bilateral supramarginal gyrus, thalamus, right superior orbital gyrus and the cerebellum ask. The r.FPN (T02) showed significant coupling with the bilateral AG, IPL, caudate and the cerebellum and decoupling in the left MFG, left supramarginal gyrus, right ATL, right IFG, SFG, precuneus, cuneus, pMTG and the cerebellum. It should be noted that the l.FPN was significantly associated with semantic processing, whereas the r.FPN was involved in pattern matching processing during the task fMRI. In the HCP data, the language task (L05) evoked significant coactivation in the IFG, SFG, MFG, thalamus, ACC, middle cingulate cortex (MCC), supplementary motor area (SMA), left supramarginal gyrus, right pMTG, ATL, cuneus, calcarine gyrus and the cerebellum and deactivation in the left IFG, mid-orbital gyrus, AG, pMTG, amygdala, hippocampus and the MOG. During WM task, the FPN (W07) was significantly synchronised with the right dorsolateral prefrontal cortex (DLPFC), IFG, ATL, pMTG, IPL, lateral occipital cortex (LOC), ITG, ACC, PCC and the cerebellum and desynchronised with the left IFG, MFG, precentral gyrus, mid-orbital gyrus and the left ITG. The motor task (M09) led to similar results to the WM task – significant coupling with the right DLPFC, IFG, ATL, MTG, LOC, ITG, IPL, ACC, PCC, precentral gyrus and the cerebellum and decoupling with the left IFG, MFG, postcentral gyrus and the left AG.

**Figure 5.**
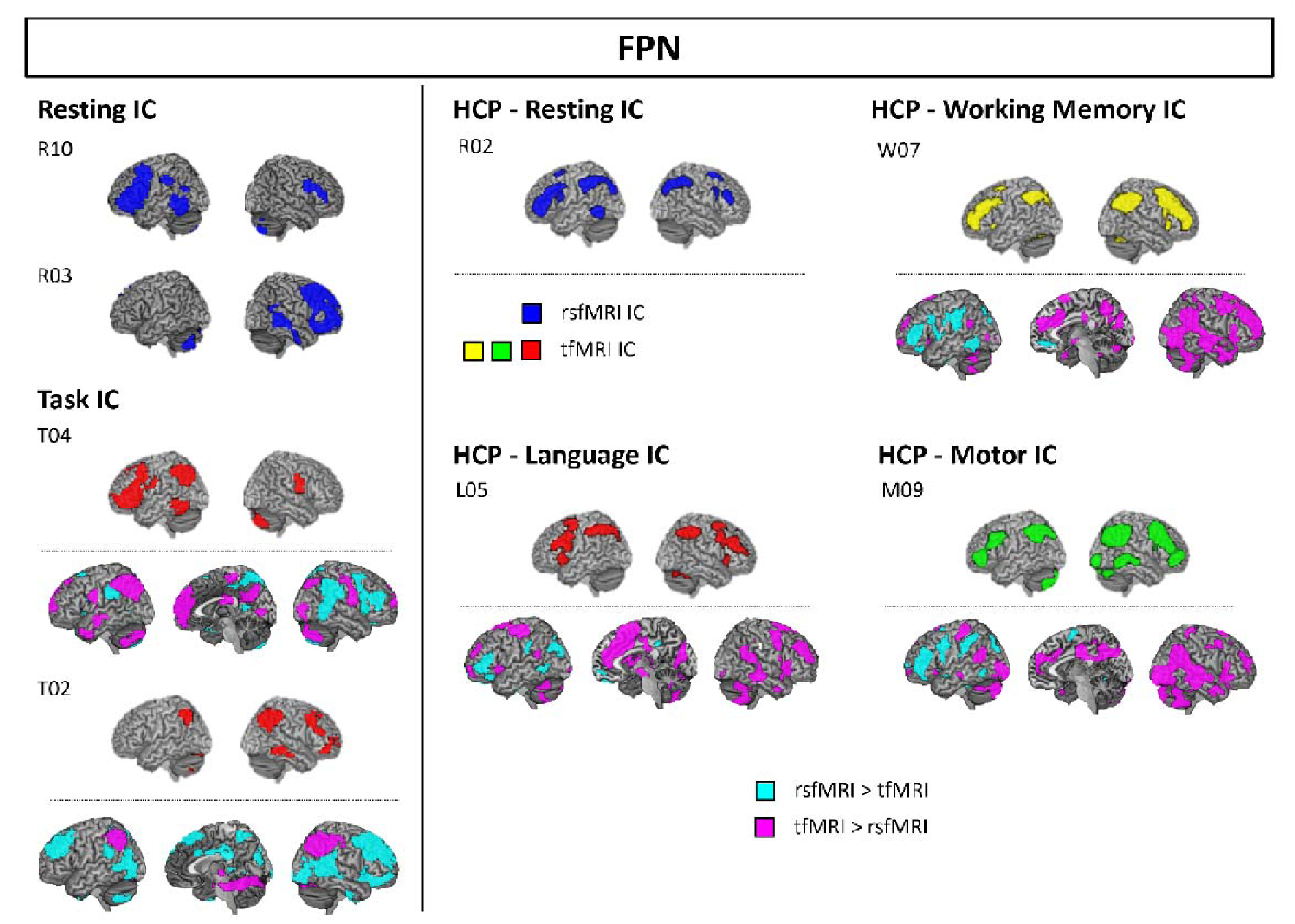
The comparison between task and rest FPN. The ICs above a dashed line are the estimated FPN from each dataset. Under the dashed line, the results of the comparison between task and rest ICs are displayed Cyan colour indicates the significant coupling in rsfMRI compared to tfMRI whereas pink colour indicates the significant coupling during tfMRI relative to rsfMRI.

The DMN revealed differential coupling and decoupling with other brain regions during task and at rest (Fig. 6). In our task ICA, there were two DMN (T14 and T28) – T14 was involved in the semantic task, whereas T28 was associated with the pattern matching task. The semantic-related DMN (T14) was more synchronised with the left DLPFC, bilateral MOG, FG, insular and MCC and desynchronised with the right IFG, PCC, pMTG, SPL and the cerebellum. The other DMN (T28) showed significant coactivation in the precuneus, IPL, SFG, precentral/postcentral gyrus, mid-orbital gyrus, IOG and cerebellum and deactivation in the ventral precuneus, MTG, and the FG. The HCP rest ICA showed anterior DMN (aDMN: R08) and posterior DMN (pDMN: R09) separately. In the language task fMRI, there was aDMN so we compared R08 with L16. The results demonstrated that the aDMN was coupled with the right IFG, supramarginal gyrus, temporal pole, mid-orbital gyrus whereas it was decoupled with the ACC, MCC, SFG, MFG, left postcentral gyrus, precuneus, midbrain, MTG and the cerebellum during the task. In the WM task fMRI, the DMN was composed of mPFC and precuneus and showed the greatest spatial similarity with the R08. We compared the WM DMN (W02) with the rest aDMN (R08). The resulted showed that the DMN was significantly coupled with the rectal gyrus, superior medial gyrus, SFG, MFG, IFG, temporal pole, precuneus and the cerebellum whereas decoupled with the bilateral SPL and the midbrain during task. In motor fMRI, the DMN (03) was matched with the rest DMN (R09) so we compared them for the analysis. The motor task evoked significant coactivation in the DLPFC, IFG, ATL, MTG, AG, mPFC, ACC, temporal pole and the cerebellum and deactivation in the precuneus, IPL, SFG, calcarine gyrus, ITG, IOG and the pMTG.

**Figure 6.**
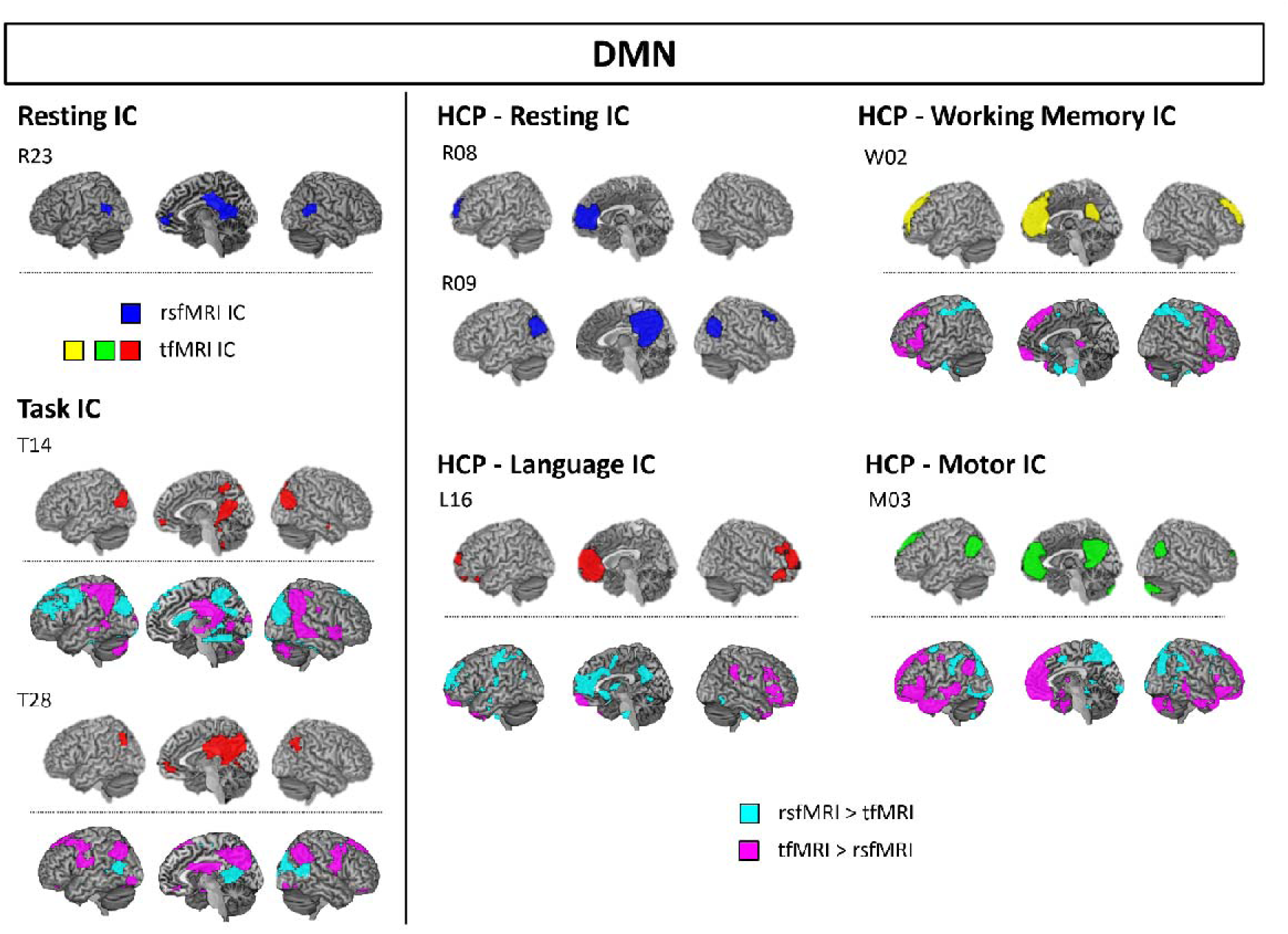
The comparison between task and rest DMN. The ICs above a dashed line are the estimated DMN from each dataset. Under the dashed line, the results of the comparison between task and rest ICs are displayed. Cyan colour indicates the significant coupling in rsfMRI compared to tfMRI whereas pink colour indicates the significant coupling during tfMRI relative to rsfMRI.

The task-specific networks showed similar results to those from the task-general networks. Tasks induced differential coupling and decoupling with other task-relevant regions in their corresponding networks (Fig. 7). The SN found in our data and HCP language data demonstrated significant coupling with the precentral gyrus, IPL, ACC, mid-orbital gyrus, PCC, FG, ITG, cuneus, thalamus and the cerebellum in our data and the temporal pole, precentral gyrus, mid-orbital gyrus and the cerebellum in HCP data during task. The SN was significantly decoupled with the ATL, IFG, DLPFC, dmPFC, precuneus, MTG, supramarginal gyrus and the right cerebellum in our data and the IPL, PCC, pMTG and the right STG in HCP data. The VN2 was more synchronised with the FG, lingual gyrus, precentral gyrus, IPL, IFG, ATL, temporal pole and the cerebellum and desynchronised with the PCC, the right postcentral gyrus and MTG during the task. The MN showed significant coactivation in the IFG, premotor cortex, SOG and the cerebellum and deactivation in the precentral/postcentral gyrus, SMA, mid-orbital gyrus, caudate, rolandic operculum, STG, MTG, ITG and the cerebellum during the task.

**Figure 7.**
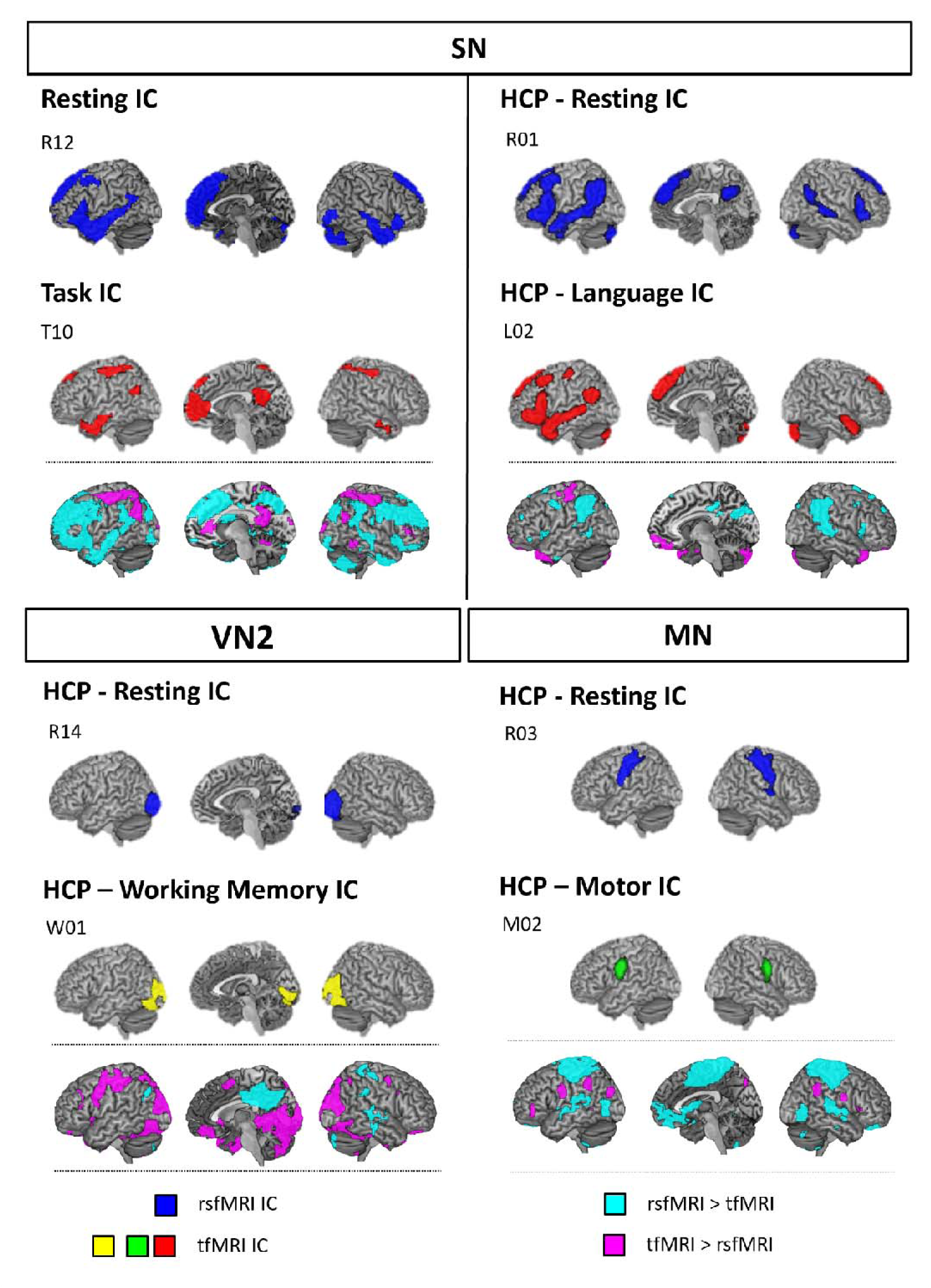
The comparison between task and rest task-specific networks. The ICs above a dashed line are the estimated networks from each dataset. Under the dashed line, the results of the comparison between task and rest ICs are displayed. Cyan colour indicates the significant coupling in rsfMRI compared to tfMRI whereas pink colour indicates the significant coupling during tfMRI relative to rsfMRI.

## Discussion

Accumulating neuroimaging evidence has shown that cognitive and motor activities are supported by collections of distributed brain regions, and these might be responsible for the underpinning cognitive computations ^35, 36^. The discovery that these distributed networks can be observed in fMRI without a task, introduced an influential notion that resting-state FC can characterize an intrinsic functional brain network and inform our understanding of cognitive function and dysfunction in patients ^3, 4^. These observations proffer the possibility that the RSNs might approximate cognitive “atomic” building blocks. Here, we examined the functional characteristics of the intrinsic and extrinsic functional network architecture by comparing rsfMRI and tfMRI. Our results demonstrated that the RSNs do not align with Dalton’s definition of atoms (indivisible, invariant and describable properties). Specifically, the task-evoked functional networks were different from the RSNs quantitatively as well as qualitatively despite some spatial correspondence between them. Furthermore, tasks reshape the RSNs by changing FC with various brain regions related to the task condition. This pattern was observed in multiple brain networks, across the task-specific networks as well as task-general networks.

Previous studies with healthy participants have focused on the overlapping spatial topology of functional networks by comparing resting and task-state networks ^9, 15, 16, 17, 18, 26^. In addition in the clinical field, there is a rapidly expanding body of research exploring functional alterations in the RSNs in neurological and psychiatric brain disorders ^37^. However, our results indicate that there is a fundamental difference in functional brain networks between rest and task conditions. We employed ICA to define the functional networks and compared the spatiotemporal properties of functional networks during task and at rest. Our results demonstrated that the functional networks estimated from the tfMRI provided more accurate task-relatedness of the networks compared to the networks from the combined data (rsfMRI and tfMRI) (Fig. 2 & Fig. S3-S5). Examining the spatial topology between rest and task-evoked networks, and consistent with previous studies, we found significant correspondence between them. Despite this, there were significant differences in the spatial similarity in the comparisons between the combined, task and rest networks – the spatial similarity between the task and rest networks was the lowest (rs ≈ 0.3) (Fig. 3). Recent studies have reported similar findings. Mennens et al ^19^ investigated the pattern of FC during 4 different tasks and at rest using a voxel-wise approach and revealed significantly reduced FC between resting and all 4 task FC (rs ≈ 0.2). Another study examined the reliability of resting and task-evoked motor network ^21^. They revealed that task fMRI showed higher correlation and better overlapping for the motor network identification than rsfMRI. The authors suggested that both tfMRI and rsfMRI could detect the motor network but rsfMRI was less reliable than tfMRI. Another study used the HCP dataset (n = 202, resting and 7 tasks) to assess various network properties between evoked and intrinsic functional networks using a graph-theory network approach ^20^. They demonstrated that despite moderate to strong overall correspondence in FC between resting and task networks, there were significant differences in both the hub structure and overall network organization between them. Furthermore, they showed a significant shift in the distribution of hub nodes from rest to task states. Taken together, these results indicate that the brain does not maintain invariant and indivisible networks when it engages in a task, and the functional interaction caused by a task can be more variable, accommodating specific task demands through flexible interactions between demand-specific regions for short periods of time ^38, 39^. Furthermore, the results indicate that caution must be taken before applying rsfMRI in clinical practice as a replacement of task fMRI – task fMRI is required to understand the full repertoire of brain functional architecture.

We used ICA to delineate functional networks, which assumes that fMRI signal from each voxel represents a linear mixture of source. It separates this signal mixture into independent source signal and groups all voxels into independent components (ICs), which represent temporally coherent functional brain networks. Several studies demonstrated that the ICA can be more sensitive to detect functional networks in tfMRI by differentiating task-related modulation in overlapping regions of two or more functional networks ^40, 41, 42, 43^. Our results revealed that ICA delineated different subsets of functional networks according to task type. Overall, ICA detected the task-general networks such as the VN, DMN and FPN across all datasets. In addition, the SN was found in our data (semantic) and the HCP language data only and the VN2 in our data and HCP WM – probably as both used pictures. Also, our tasks (semantic and picture matching) separated the FPN across the hemispheres: l.FPN was involved in semantic processing whereas r.FPN was coupled with the picture matching task. The DMN showed differential configuration according to the dataset. In our data, we found one DMN in rsfMRI and two DMN in tfMRI serving each task condition (T14: semantic; T28: picture matching). In HCP data, there were two sets of the DMN at rest (aDMN and pDMN) and one in each tfMRI. Our results indicate that cognitive function is supported by the interaction between distributed brain regions and tasks require dynamic reconfiguration of the functional brain networks ^38, 39^. A very similar outcome is found in large PDP neurocomputational models of language and other functions; where centres and pathways show changing divisions of labour both across contrastive tasks (e.g., repetition vs. comprehension) but also within a task corresponding to the characteristics of specific words/stimuli ^44, 45^. Also, our findings add considerable support to the conclusion that the FPN and DMN are dynamic and consist of several subsystems, serving distinct cognitive functions (FPN: language, attention, working memory, memory, spatial cognition, action, sensation, and perception; DMN: semantic memory, episodic memory, decision making, and social cognition) ^46, 47, 48, 49^. Furthermore, data-driven multivariate approaches can provide further insights into differences in brain network across varying conditions that go beyond simple comparisons of the spatial correlation in FC between them.

In conclusion, our findings suggest that task-based functional MRI is essential for gaining a comprehensive understanding of the brain’s functional architecture. This approach provides insights into how the brain dynamically adjusts its network connectivity to meet specific task demands, demonstrating the need to consider both resting-state and task-based fMRI for a comprehensive perspective on brain function.

## Acknowledgements

The research was supported by the AMS Springboard (SBF007\100077) to J.J and a MRC Programme Grant (MR/R023883/1) and MRC intramural funding (MC_UU_00005/18) to M.A.L.R.

