## Supplementary material for "Are resting-state networks the brain’s cognitive atoms? Differential dynamic reconfiguration of functional brain networks across tasks and at rest": SI

**Supplementary information**

Supplementary figure 1

Supplementary figure 2

Supplementary figure 3

Supplementary figure 4

Supplementary figure 5

Supplementary figure 6

Supplementary table 1

Supplementary table 2

**Supplementary Figure 1**

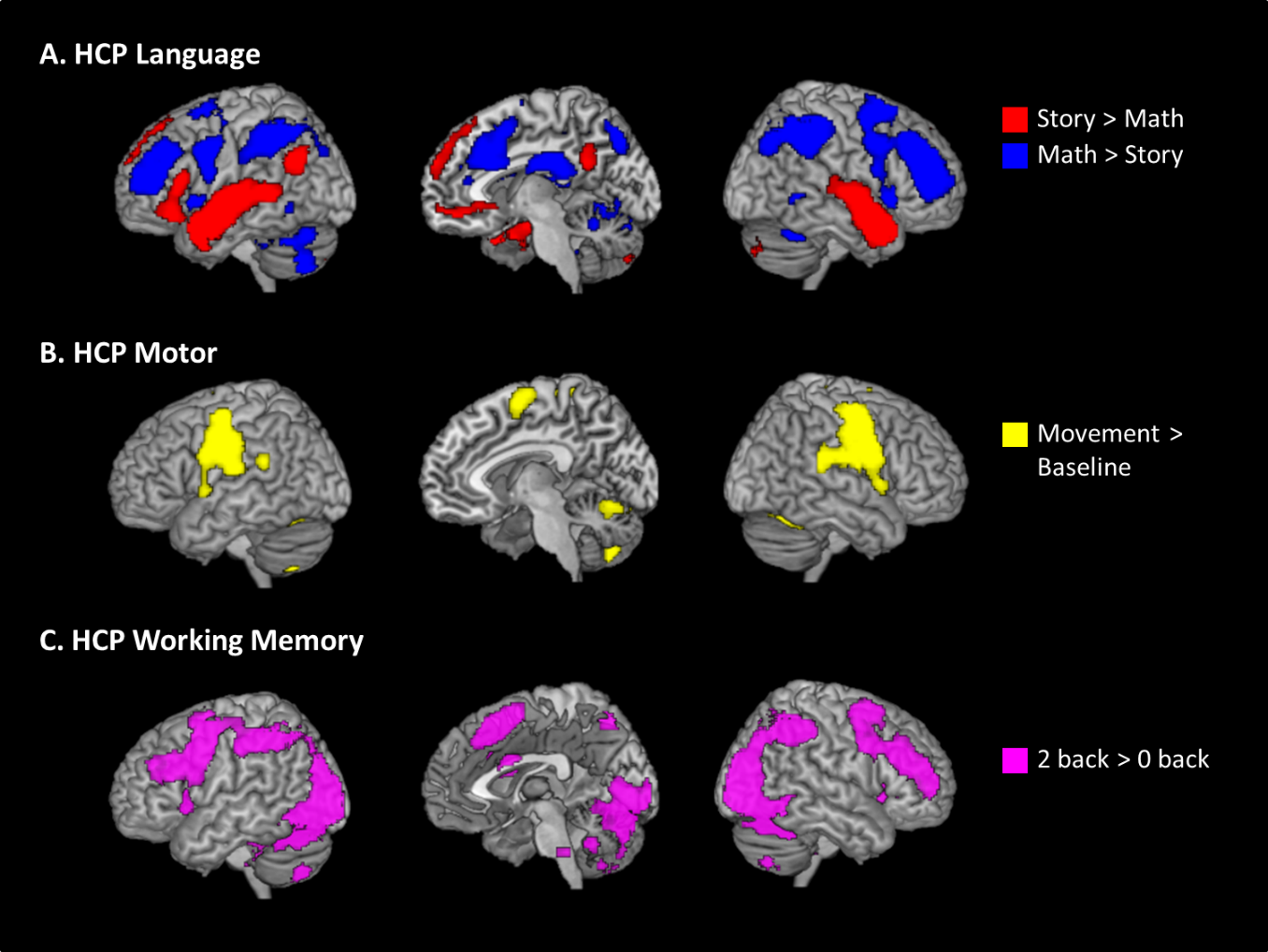

Figure S1. The GLM results of HCP data. Statistical threshold was set at p < 0.001 at the voxel-level and p < 0.05 at the cluster level with at least 100 contiguous voxels after family-wise error (FWE) correction.

**Supplementary Figure 2**

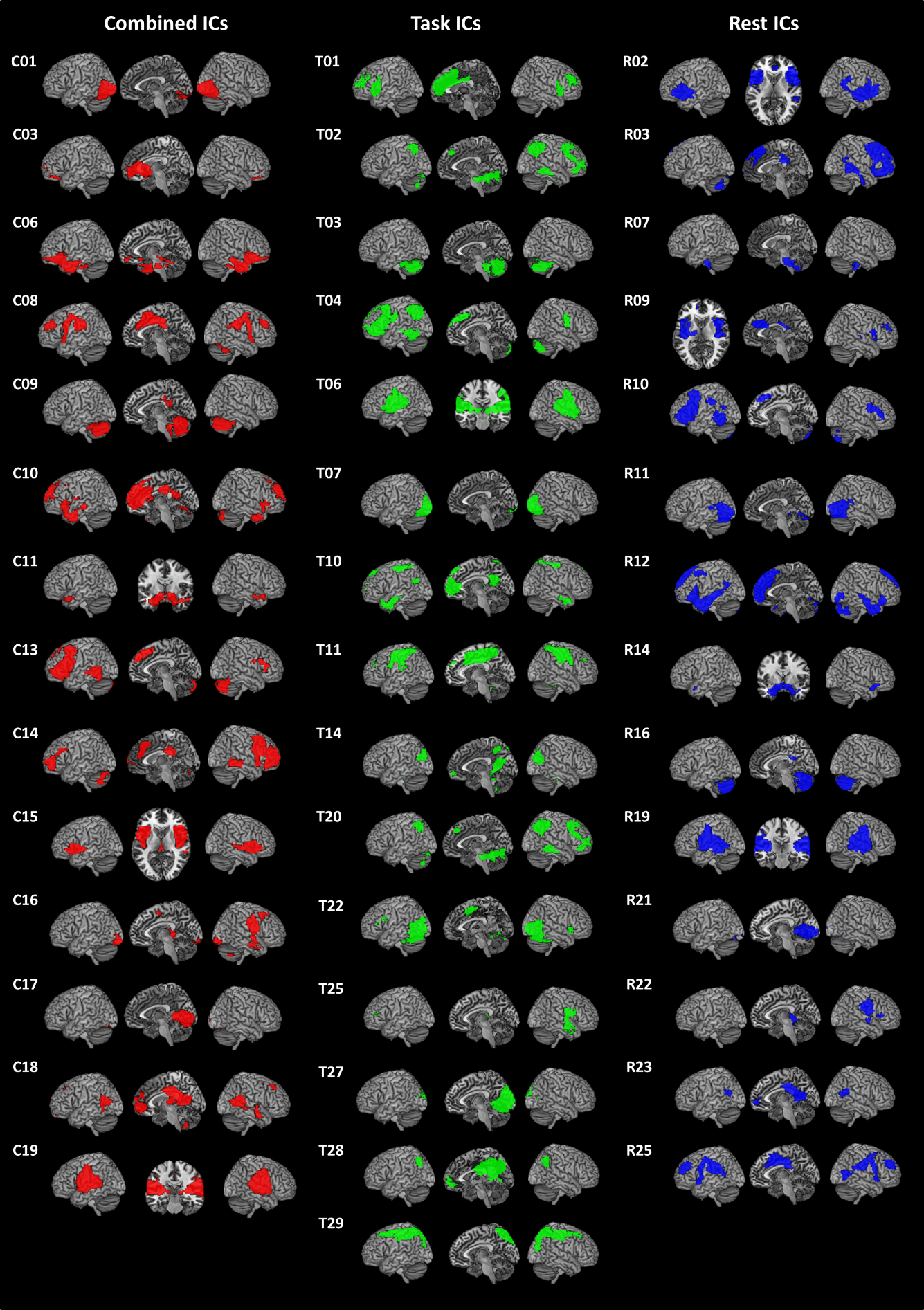

**Figure S2.** The results of 3 ICAs.

**Supplementary Figure 3**

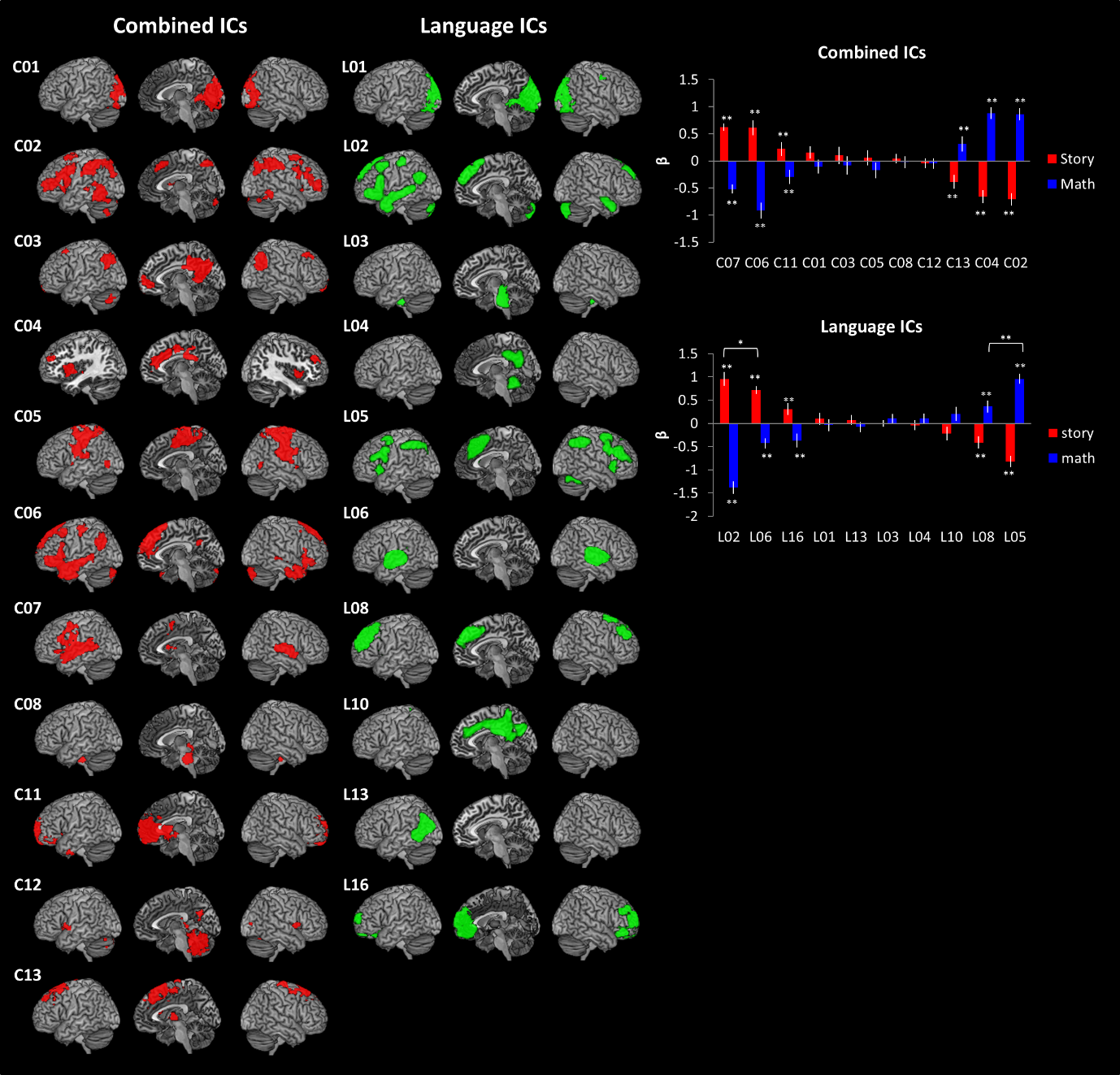

**Figure S3.** The results of HCP Language. * p < 0.05, ** p < 0.005.

**Supplementary Figure 4**

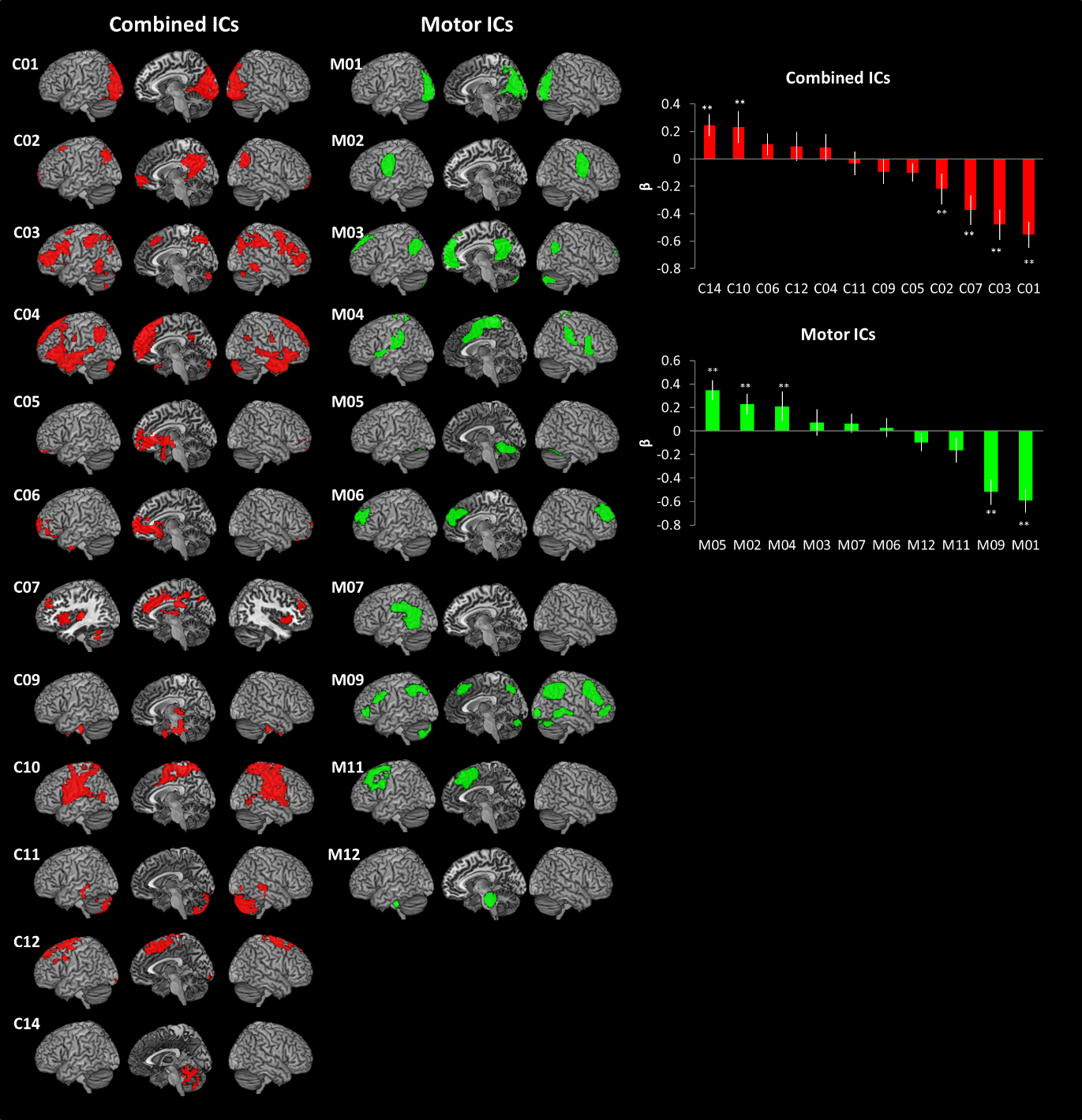

**Figure S4.** The results of HCP Motor. ** p < 0.005.

**Supplementary Figure 5**

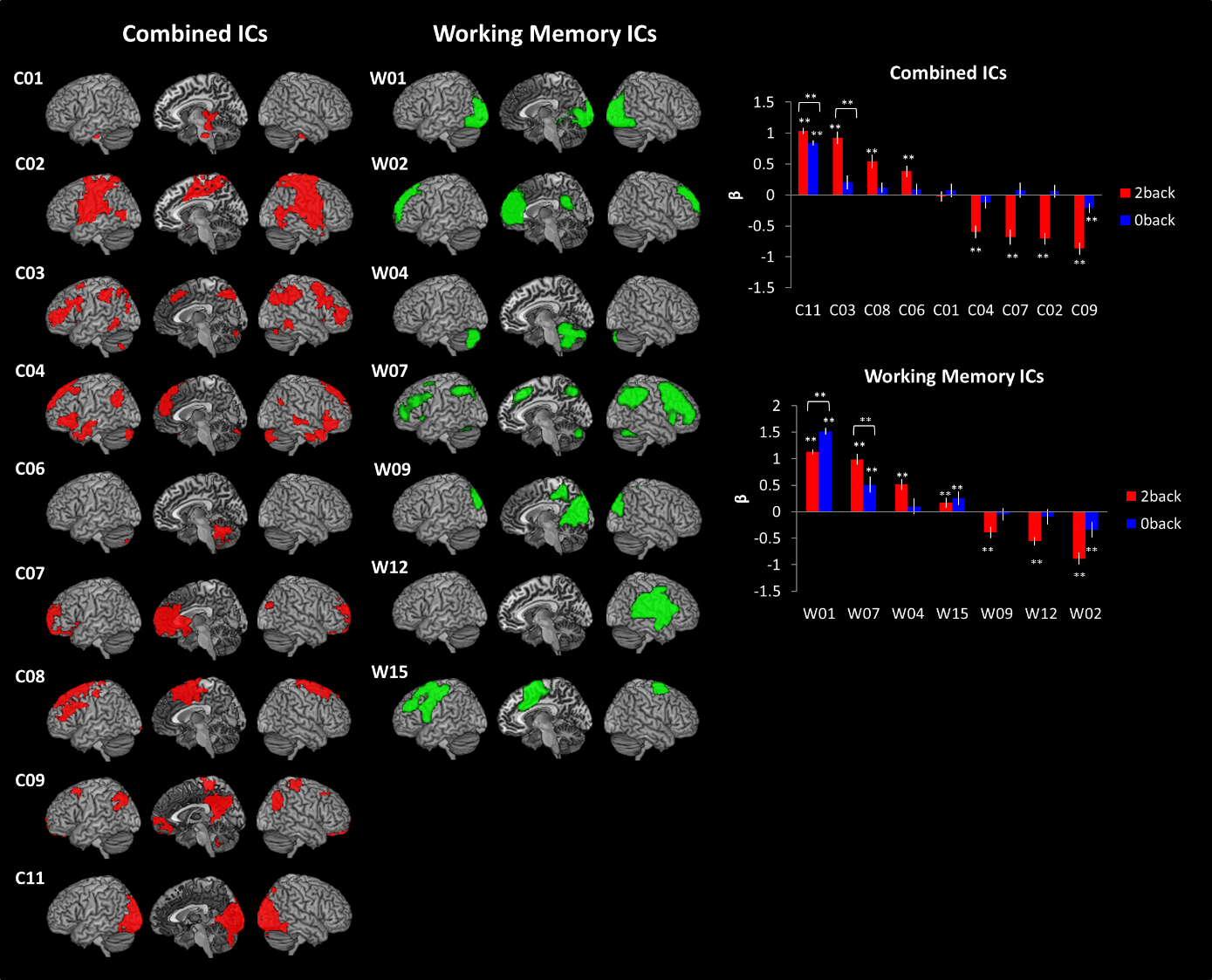

**Figure S5.** The results of HCP Working Memory. ** p < 0.005.

**Supplementary Figure 6**

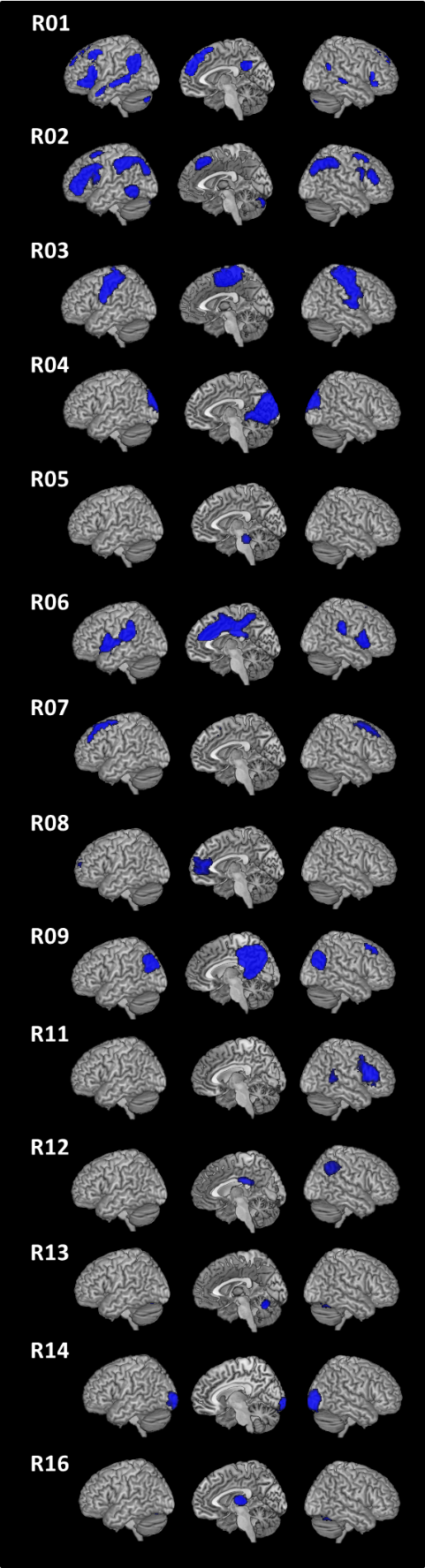

**Figure S6.** The results of HCP resting-state fMRI.

**Supplementary Table 1**

|  | IC | Semantic | Control |
| --- | --- | --- | --- |
| Combined ICA | C01 | 0.05 | -0.04 |
|  | C03 | -0.15 | 0.12 |
|  | C06 | 0.02 | -0.11 |
|  | C08 | -0.07 | 0.03 |
|  | C09 | 0.07 | -0.04 |
|  | C10 | 0.02 | 0.01 |
|  | C11 | 0.03 | -0.09 |
|  | C13 | -0.10 | 0.00 |
|  | C14 | -0.02 | 0.09 |
|  | C15 | -0.10 | 0.04 |
|  | C16 | -0.01 | -0.08 |
|  | C17 | 0.05 | -0.03 |
|  | C18 | 0.06 | -0.13 |
|  | C19 | 0.00 | 0.02 |
| Task ICA | T01 | 0.02 | 0.02 |
|  | T02 | **-0.38** | **0.46** |
|  | T03 | -0.11 | **0.43** |
|  | T04 | **1.08** | **-0.93** |
|  | T06 | **-0.34** | **0.20** |
|  | T07 | **0.34** | **0.48** |
|  | T10 | **0.49** | **-0.81** |
|  | T11 | -0.12 | **0.18** |
|  | T14 | **0.44** | **-0.42** |
|  | T20 | **-0.17** | **0.17** |
|  | T22 | **1.34** | **-0.82** |
|  | T25 | **-0.14** | **0.16** |
|  | T27 | -0.17 | **0.49** |
|  | T28 | **-0.76** | **0.40** |
|  | T29 | **-0.60** | **0.94** |

**Table S1.** The results of temporal regression. The value indicates the mean of β resulted from multiple regression. Bold indicate the ICs significantly related to task conditions (p _FDR-corrected_ < 0.005).

**Supplementary Table 2**

|  | Language | | | Motor | | Working Memory | | |
| --- | --- | --- | --- | --- | --- | --- | --- | --- |
|  | IC | Story | Math | IC | Movement | IC | 0 Back | 2 Back |
| Combined ICA | C01 | 0.15 | -0.11 | C01 | **-0.55** | C01 | 0.07 | -0.03 |
|  | C02 | **-0.71** | **0.86** | C02 | **-0.22** | C02 | 0.07 | **-0.71** |
|  | C03 | 0.10 | -0.08 | C03 | **-0.48** | C03 | 0.20 | **0.93** |
|  | C04 | **-0.66** | **0.88** | C04 | 0.09 | C04 | -0.12 | **-0.59** |
|  | C05 | 0.06 | -0.18 | C05 | -0.10 | C06 | 0.09 | **0.38** |
|  | C06 | **0.61** | **-0.92** | C06 | 0.11 | C07 | 0.08 | **-0.68** |
|  | C07 | **0.63** | **-0.52** | C07 | **-0.37** | C08 | 0.12 | **0.54** |
|  | C08 | 0.04 | -0.03 | C09 | -0.09 | C09 | **-0.21** | **-0.87** |
|  | C11 | **0.22** | **-0.30** | C10 | **0.23** | C11 | **0.84** | **1.04** |
|  | C12 | -0.04 | -0.05 | C11 | -0.03 |  |  |  |
|  | C13 | **-0.39** | **0.31** | C12 | 0.09 |  |  |  |
|  |  |  |  | C14 | **0.25** |  |  |  |
| Task ICA | L01 | 0.10 | -0.04 | M01 | **-0.59** | W01 | **1.52** | **1.13** |
|  | L02 | **0.96** | **-1.38** | M02 | **0.23** | W02 | **-0.35** | **-0.89** |
|  | L03 | 0.00 | 0.11 | M03 | 0.07 | W04 | 0.10 | **0.51** |
|  | L04 | -0.04 | 0.11 | M04 | **0.21** | W07 | **0.51** | **0.99** |
|  | L05 | **-0.82** | **0.96** | M05 | **0.35** | W09 | -0.05 | **-0.39** |
|  | L06 | **0.72** | **-0.43** | M06 | 0.03 | W12 | -0.10 | **-0.56** |
|  | L08 | **-0.41** | **0.36** | M07 | 0.07 | W15 | **0.25** | **0.17** |
|  | L10 | -0.22 | 0.20 | M09 | **-0.52** |  |  |  |
|  | L13 | 0.07 | -0.08 | M11 | -0.16 |  |  |  |
|  | L16 | **0.31** | **-0.37** | M12 | -0.10 |  |  |  |

**Table S2.** The results of temporal regression from HCP data. The value indicates the mean of β resulted from multiple regression. Bold indicate the ICs significantly related to task conditions (p _FDR-corrected_ < 0.005).
